# Anyone can be the best: Impact of diverse methodologies on the evaluation of structural variant callers

**DOI:** 10.1101/2025.08.28.672546

**Authors:** Luca Denti, Thomas Krannich, Tomas Vinar, Rayan Chikhi, Paola Bonizzoni, Brona Brejova, Fereydoun Hormozdiari

## Abstract

Structural variants (SVs) are medium and large-scale genomic alterations that shape phenotypic diversity and disease risk. Numerous methods have been proposed for discovering SVs, however their benchmarking has been inconsistent across studies, often resulting in contradictory findings. One of the main sources of conflicting evaluation results is the lack of consistency in the SV callsets used as ground truth, ranging from curated callsets released by consortia to more recent approaches that construct callsets from high-quality telomere-to-telomere *de novo* haplotype assemblies. The discrepancies between benchmarks are further compounded by the choice of the reference genome (`GRCh37`, `GRCh38`, and `T2T-CHM13`), where using `T2T-CHM13`reveals a different deletion/insertion profile, indicating reduced reference bias. We evaluated the performance of several state-of-the-art SV discovery methods from long-read whole-genome sequencing data and observed substantial variation in their performance and rankings, depending on the choice of ground truth, reference genome, and genomic regions used for evaluation. Counter-intuitively, the more complete reference genome `T2T-CHM13`does not inherently solve the problem of SV benchmarking; instead it reveals the limitations of each detection method in complex genomic regions. The substantial variation in detection accuracy across different genomic regions calls for additional caution in downstream analyses and in drawing conclusions based on predicted SVs. These findings underscore the complexity of evaluating SV detection methods and highlight the need for careful consideration and, ideally, field-standard best practices when reporting performance metrics.

## 1 Introduction

Genomic structural variants (SVs) are traditionally defined as medium and large-scale (*>*50bp) genomic alterations, including deletions, duplications, inversions, translocations, and more complex rearrangements [1,2,3]. It has become evident that SVs play a profound role in shaping phenotypic diversity and disease susceptibility [3]. The advent of highthroughput sequencing technologies and sophisticated computational methods has revolutionized our ability to detect and characterize these previously elusive SVs, revealing their pervasive nature and immense impact on human health [4,5,6,7,8,9]. Historically, short-read sequencing platforms, such as Illumina, have been the workhorse of genomics due to their high throughput and low cost per base. While effective for detecting smaller and simpler SVs, short reads often struggle to resolve complex rearrangements, variants in repetitive regions, or large insertions due to their limited read length, leading to challenges in breakpoint resolution and an increased rate of false positives. Furthermore, recent analyses have indicated that a substantial fraction of SVs can not be detected using short-read WGS data [10,11].

The emergence of long-read sequencing technologies, including Pacific Biosciences (PacBio) and Oxford Nanopore Technologies (ONT), has revolutionized SV detection [3,12]. Reads that are tens of kilobases long can span entire SVs, enabling direct detection of breakpoints, precise characterization of variant architecture, and improved phasing of SVs with other genetic variations, even in complex and highly repetitive genomic regions. Over the past few years, many tools have been developed for SV prediction from long-read WGS data (Table 1), along with growing number of comparative evaluations [13,12,14]. SV detection typically starts with alignment to a reference genome using tools like `minimap2`[15], detecting signatures such as large gaps, large insertions, or split alignments. Subsequent steps often involve clustering these individual signatures to form consensus calls, refining breakpoints to base-pair precision, and genotyping the identified variants. State-of-the-art long-read SV callers typically use advanced algorithmic strategies, including graph-based approaches, or machine learning models [3].

**Table 1:** State-of-the-art methods for SV discovery from long reads WGS data and their comparative evaluation settings. CMRG stands for Challenging Medically Relevant Genes, the recent SV callsets curated by the GIAB consortium [17]. `benchmark_callset.py`is a `python`script available at https://github.com/Maggi-Chen/DB_code

| Tool | Reference |  |  | Ground Truth |  |  |  | Benchmarking Tool |
| --- | --- | --- | --- | --- | --- | --- | --- | --- |
|  | GRCh37 | GRCh38 | T2T-CHM13 | GIAB v0.6 | GIAB v1.1 | CMRG | assembly-based |  |
| [28] cuteSV (2020) | ✓ |  |  | ✓ |  |  |  | truvari |
| [16] SVDSS (2022) | ✓ | ✓ |  | ✓ |  | ✓ | dipcall | truvari |
| [29] SVision (2022) | ✓ |  |  | ✓ |  |  |  | truvari |
| [21] sniffles2 (2023) | ✓ | ✓ |  | ✓ |  | ✓ | dipcall | truvari |
| [22] debreak (2023) | ✓ |  | ✓ | ✓ |  |  | paf, dipcall | benchmark_callset.py |
| [30] SVision-pro (2023) |  | ✓ |  | ✓ |  |  |  | truvari |
| [23] Severus (2025) | ✓ | ✓ |  | ✓ |  |  | hapdiff | truvari, minda |
| [25] sawfish (2025) |  | ✓ | ✓ |  | ✓ | ✓ |  | truvari refine, hap-eval |

While long-read sequencing greatly enhances SV detection, accurately identifying SVs in certain genomic regions remains challenging [16,17]. Factors such as read misalignment in repetitive regions, incorrect clustering of gaps and insertions between alignments, sequencing errors, and low coverage can contribute to both false positives and missed variants. Moreover, complex SVs, closely spaced or overlapping variants on different haplotypes, and multi-breakpoint events are often misclassified, fragmented, or incorrectly merged by existing tools, resulting in inaccurate SV calls. In addition, the best practices and optimal evaluation strategies remain undefined, and benchmarking results often vary across studies, leaving the current performance assessment landscape inconsistent and difficult to interpret. More importantly, this has created ambiguity about the true capabilities of leading methods to accurately detect SVs, with far-reaching implications for routine genome analysis. SV detection methods are typically evaluated against curated callsets, such as those from the Genome in a Bottle (GIAB) consortium [18,17], or against callsets identified from available high-quality Telomere-to-Telomere (T2T) *de novo* assemblies using tools like `dipcall`[19] or `hapdiff`[20]. The latter approach is gaining popularity as leveraging T2T *de novo* assemblies has proven exceptionally powerful, as it enables detection of SVs in repetitive regions of the genome and allows for a more comprehensive and accurate evaluation of SV detection tools [16, 21, 22, 23, 24].

Although most studies overall follow similar high-level evaluation strategies, performance evaluations have produced conflicting and inconsistent results, with some studies reporting F1 scores above 0.95 for SV detection [25] while others suggest that current state-of-the-art tools achieve substantially lower performance, with F1 scores as low as 0.7 [26]. We believe that these performance discrepancies arise mainly from the distinct settings used for each evaluation. To prove our claim, in this work, we investigate the choices that can affect the evaluation of SV callers, e.g., which reference genome is used, which regions of the genome are considered, which ground truth is considered, and which benchmarker tool is used. Resolving these discrepancies is crucial, as these inconsistencies not only affect how biologists analyze long-read whole-genome sequencing (WGS) data but also obscure the true strengths and limitations of current SV detection methods. This uncertainty, in turn, compromises the accuracy and reliability of biological interpretations derived from SV predictions in downstream analyses.

## 2 Results

This study investigates the performance disparities among state-of-the-art methods for SV discovery, addressing existing inconsistencies in reported SV detection results. Our analysis leveraged the well-characterized HG002 sample, utilizing its comprehensive resources: long-read HiFi PacBio data, high-quality *de novo* assembly, and the GIAB SV callsets (v0.6 and v1.1). This sample and and its available high-quality *de novo* assembly and curated SVs callsets (such as GIAB v0.6) have been widely used in performance evaluation of SV callers. Table 1 summarizes the various recent methods published for SV discovery from long read WGS data, along with the reference genomes and ground truths used in their evaluations.

Through international consortium efforts such as the GIAB project, comprehensive SV callsets have been curated for a select number of extensively studied genome samples. These callsets are widely recognized as the gold standard for SV benchmarking, with the majority of SV callers having been evaluated against them (see Table 1). Within the scope of this manuscript, we will refer to SV callsets provided by such consortiums (including the GIAB v0.6 and v1.1 callsets) as *curated SV callsets*. Furthermore, the emergence of high-quality T2T (or near-T2T) phased *de novo* genome assemblies has opened new avenues for generating robust SV callsets. Tools such as `dipcall`[19], `SVIM-asm`[27], and `hapdiff`[20] leverage these assemblies to produce high-confidence SV calls, which have subsequently been utilized for benchmarking various SV methodologies (see Table 1). Throughout this paper, we will designate SVs derived from high-quality T2T (or nearT2T) *de novo* assemblies as *assembly-based SV callsets*. In contrast, SV callsets generated directly from raw sequencing reads using various SV prediction methods (Table 1) will be referred to as *read-based SV callsets*.

### 2.1 Discrepancies in assembly-based SV callsets

The most reliable approach for constructing robust and comprehensive SV callsets for any sample is based on aligning high-quality *de novo* assemblies to the reference genome of interest (e.g., `GRCh38`or `T2T-CHM13`) [16,21,22,23,24]. This is also the methodology adopted by the GIAB consortium to curate the recent SV callset on Challenging Medically Relevant Genes (CMRG) [17] and the preliminary HG002 v1.1 callset (unpublished). We applied this strategy to construct assembly-based SV callsets for the HG002 sample using the high-quality *de novo* assembly provided by the Human Pangenome Reference Consortium (HPRC) [31] and three leading assembly-based SV callers: `dipcall`[19], `SVIM-asm`[27], and `hapdiff`[20]. Note that `dipcall`and `hapdiff`have been adopted for evaluating SV prediction methods in many of the recent publications (Table 1). We also decided to include `SVIM-asm`since `hapdiff`is based on a modified version of this approach. The three state-of-the-art assembly-based SV calling tools were run on the same highquality *de novo* HG002 assembly and assembly-based SVs callsets were generated against three major human reference genomes: `GRCh37`, `GRCh38`, and `T2T-CHM13`. We restricted our analyses on these callsets to variants larger than 50bp, with called genotype different from homozygous reference (i.e., `0|0`), and falling in those regions reported as *confident* by the callers. We observed substantial differences in the properties of the generated assembly-based SV callsets, as well as distinct biases associated with each callset, as detailed below. Figure 1 reports the results of this analysis. Surprisingly, when considering the full genome (i.e., not restricting our analysis to confident regions), we observed a similar trend (Supplementary Figure S1).

**Fig. 1:**
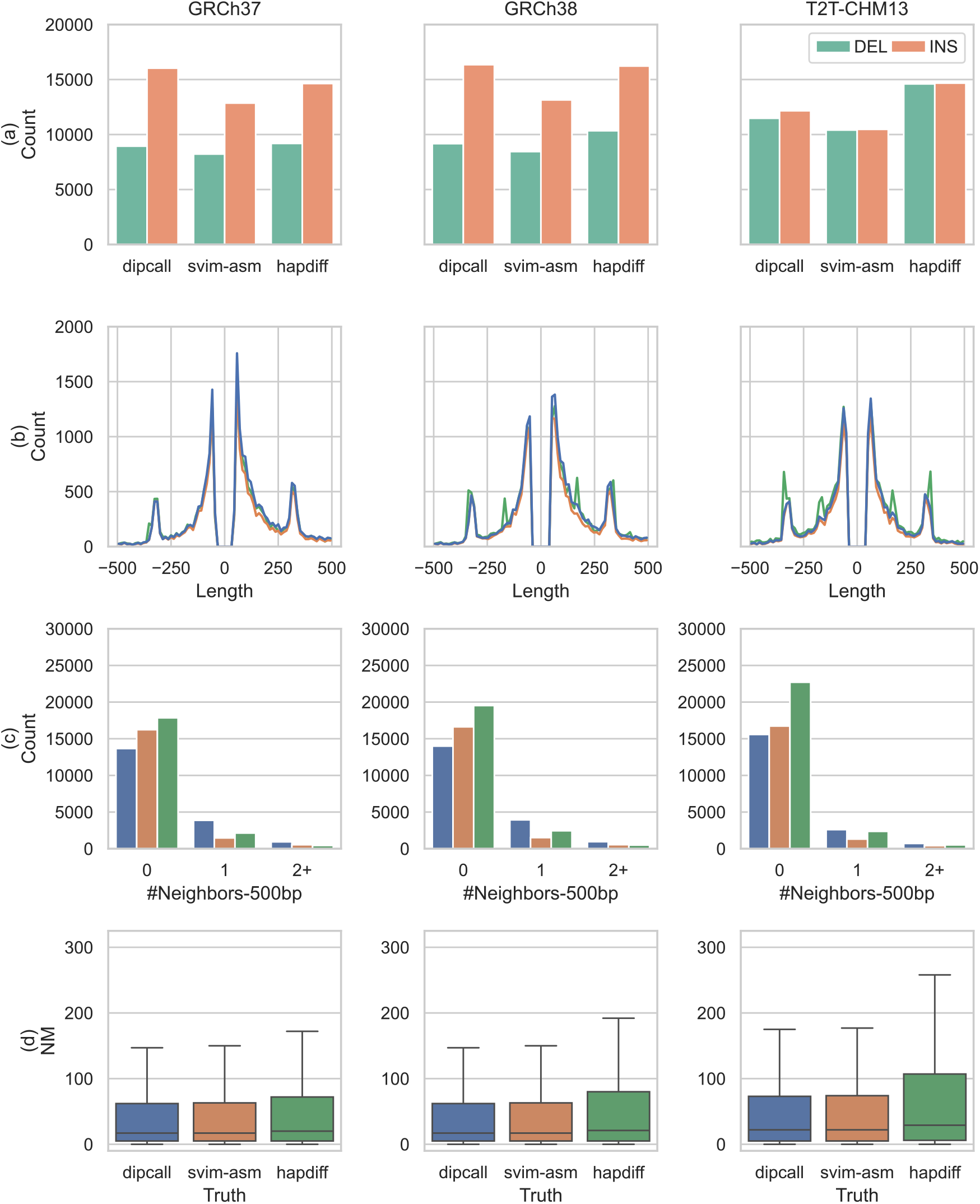
Comparison of state-of-the-art assembly-based SV callers using the same *de novo* assembly against three reference genomes. (a) Number of SVs reported for each approach based on their type (INSertions and DELetions). (b) Length distribution of unbalanced structural variants (negative length for deletions). (c) Number of neighboring SVs within 500bp. (d) Distribution of the total number of mismatches and gaps (`NM`field) when aligning “alternative” contigs created from the VCF produced by each method against the true contigs. Results on full genome are presented in Supplementary Figure S1.

### Number and length of SVs

First, we observed substantial differences in the total number of SVs identified across the assembly-based SV callsets generated by the three state-of-the-art methods (`dipcall`, `SVIM-asm`, and `hapdiff`), despite all using the same input data (Figure 1(a)). Furthermore, when `GRCh38`or `GRCh37`is used as the reference genome, all three methods exhibit a consistent bias toward detecting more insertions than deletions. In contrast, using the `T2T-CHM13`reference yields a more balanced representation of insertions and deletions across all three callers (Figure 1(a)). In addition, we also observed considerable differences in the length distribution of the identified SVs based on both the method and the reference genome utilized (Figure 1(b)). The observed differences in SV length likely reflect biases in the ability of each method to accurately detect specific SV types, especially in low complexity regions where SV calling is more sensitive to alignment parameters and calling heuristics.

### Distribution of closely spaced SVs

One major source of complexity in SV detection and evaluation is the close proximity of SVs within certain regions of the genome, either on the same haplotype or across different haplotypes. To assess this, we quantified the number of SVs identified by each assembly-based SV detection tools that are located within 500bp of each other (Figure 1(c)). They were stratified by the method used for the SV callset construction (`dipcall`, `SVIM-asm`, and `hapdiff`) and reference genome used (`T2T-CHM13`, `GRCh38`, and `GRCh37`). We observed substantial differences across the three methods, with `dipcall`in particular showing increased frequency of reported closely spaced SVs. On `GRCh37`and `GRCh38`, 26% of `dipcall`calls are closely spaced against 13% and 11% of `hapdiff`and `SVIM-asm`. These percentages drop to 17%, 11%, and 9% when considering the `T2T-CHM13`reference genome. This suggests that `dipcall`may be more prone to identifying clustered SVs, which could either reflect true biological complexity or method-specific artifacts.

### Comparison of SV calls to original contigs

Inspired by the methodology proposed in [32], we investigated the agreement between each assembly-based SV callset and the *de novo* assembly from which the corresponding callset was derived. For each SV callset generated by the three approaches, we constructed alternative contigs by integrating the predicted SVs into the reference genome and extracting 500bp of flanking sequence on each side (SVs located within 500bp of one another were merged into a single contig). The resulting alternative contigs were then aligned back to the original HG002 *de novo* assembly with `minimap2`[15]. We then considered all primary alignments and assessed their accuracy based on the total number of mismatches and gaps, as reported in the `NM`field (Figure 1(d)). Based on this analysis we observed the lowest mismatches and gaps (measured by the median of `NM`field) in alternative contigs generated from `dipcall`and `SVIM-asm`followed by `hapdiff`(Figure 1(d)). We recall that both `dipcall`and `SVIM-asm`share the same alignments and same confident regions, thus similar results are expected. However, we note that `dipcall`contigs exhibit a slightly lower upper quartile (1 point). Notably, the median of `NM`increased across all methods as the completeness of the reference genome improved. On `GRCh37`reference genome, the median `NM`values were 17 and 20 for alternative contigs generated by SV predictions from `dipcall`/`SVIM-asm`and `hapdiff`, respectively. On `GRCh38`reference genome, the median `NM`values remain the same for `dipcall`/`hapdiff`and increase to 21 for `hapdiff`whereas on `T2T-CHM13`reference genome, they increased to 22 and 29. This trend is likely due to the inclusion of more complex genomic regions in the latest `T2T-CHM13`reference genome. Note that the alternative contigs do not integrate any SNPs or small indels, so some variation in the alignments to the *de novo* assembly is expected.

### Assessment of divergence between assembly-based SV callsets

Finally, we assessed the overall agreement between the assembly-based SV callsets generated by each method from the same high-quality *de novo* assembly. To do this, we compared each pair of SV callsets using `truvari`[33], a widely used SV benchmarking toolkit. When considering the SV falling in confident regions, we observed some degree of divergence between the three SV callsets, even when SV representations were harmonized using `truvari refine`(Figure 2). Pairwise similarity scores between assembly-based SV callsets ranged from as high as 0.95 (`dipcall`vs `SVIM-asm`on `GRCh37`and `GRCh38`) to as low as 0.77 (`hapdiff`vs `SVIM-asm`on `T2T-CHM13`), indicating method-specific differences in assembly-based SV callset construction. As expected, since they use the same alignments and confident regions, `dipcall`and `SVIM-asm`show higher similarity, while their similarity to `hapdiff`is consistently lower. Notably, the similarity between `dipcall`and `hapdiff`never exceeds 0.87 in any setting we considered, and falls below 0.5 when considering the full genome (Supplementary Figure S2). Even more concerning, we observed that the overall similarity among assembly-based SV callsets generated by the three methods decreases as the completeness of the reference genome increases (Figure 2 and Supplementary Figure S2). In particular, the greatest divergence between methods was observed when using `T2T-CHM13`as the reference genome. This suggests that more complete references might either expose method-specific biases or introduce greater detection complexity for specific regions of the genome. Similarity between callsets is even lower when SV representations are not harmonized, i.e., when similarity is computed using `truvari bench`(Supplementary Fig-S2a ures and S2c). A similar trend, albeit with more pronounced differences between the assembly-based SV callsets, is still observed when the analysis is not limited to confident S2a regions of the genome (Supplementary Figures and S2b). Notably, when focusing on full genome, the impact of using different reference genomes increases. This result further supports the notion that incorporating more complex, low-mappability regions in a complete reference amplifies the discrepancies among state-of-the-art assembly-based SV callers.

**Fig. 2:**
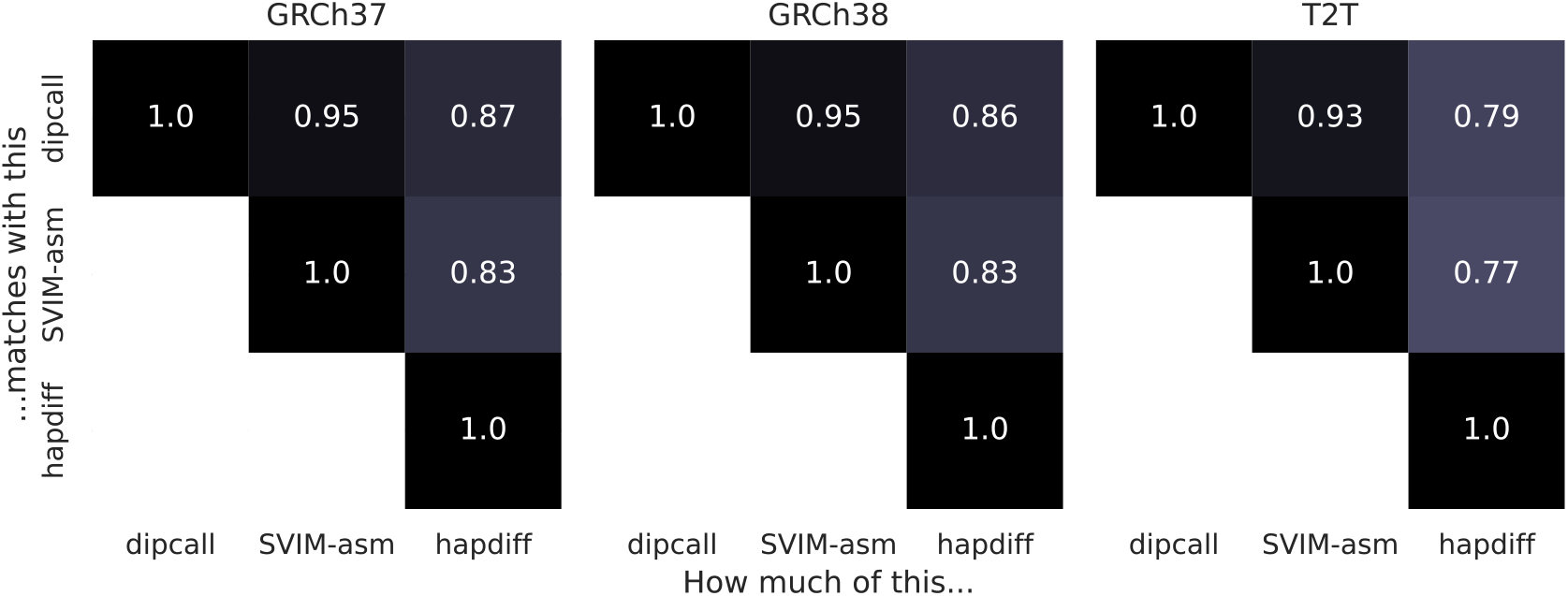
Pairways similarity between assembly-based SV callers. Each cell presents the accuracy (TP/(TP+FP+FN)) of the column tool (considered as the callset) w.r.t. the row tool (considered as the truth set). Each cell reports the accuracy on the full genome, computed using `truvari refine`. The same analysis on full genome without using `truvari refine`and on confident regions reported by the various tools is presented in Supplementary Figure S2.

### 2.2 Effect of Ground Truth and Reference Selection on Read-Based SV Evaluation

Benchmarking newly developed SV detection methods against established ground truths is standard practice, but the observed discrepancies in reported evaluation results across comparative studies raise critical questions and create ambiguity about the true capabilities of such methods. In this section, we explore the sources of these inconsistencies and we quantify the impact of user-defined factors, such as the choice of the ground truth, on the observed performance metrics and rankings in these comparative evaluations. Our results suggest that a major source of variability in the evaluation of SV prediction tools stems from the use of different ground truths and reference genomes (e.g., `GRCh37`, `GRCh38`, and `T2T-CHM13`). We will investigate the impact of diverse SV callsets commonly used as ground truths in the evaluation of read-based SV callsets: assembly-based SV callsets and curated SV callsets.

We considered the HG002 PacBio HiFi dataset (Sequel II System with Chemistry 2.0, 15kb Library, 36x coverage) provided by the Human Pangenome Reference Consortium (HPRC) [31] and ran seven long reads-based SV prediction callers: `cuteSV`, `SVDSS`, `sniffles2`, `debreak`, `SVision-pro`, `Severus`, and `sawfish`.

### Benchmarking of read-based SV callsets against assembly-based ground truths

To evaluate performance, we compared each read-based SV callset against three assembly-based callsets serving as ground truths. These callsets were generated from the highquality *de novo* assembly provided by the HPRC using `dipcall`, `SVIM-asm`, and `hapdiff`.

This evaluation analysis was conducted across the three primary human reference genomes: `GRCh37`, `GRCh38`, and `T2T-CHM13`. Performance of each SV caller was quantified using `truvari`under three distinct parameter settings, denoted as `truvari-def`, `truvari-wbed`to assess robustness (see Section 3 for details). We observed that the F1 score of each tool varies substantially depending on the choice of ground truth, reference genome, and `truvari`parameters. Figure 3a reports the results stratified by reference genome (full results are provided in Supplementary Table 1). On average, all read-based SV callers exhibit lower accuracy when using the recent `T2T-CHM13`reference genome. When considering the full genome and SV harmonization, the average F1 score drops from 81% on `GRCh37`and 77% on `GRCh38`to 62% on `T2T-CHM13`. Accuracy of all tools improves when restricting the analysis to confident regions of the genome (as reported by the assembly-based callers). In this setting, the average F1 score increased to 85%, 84%, and 74% on `GRCh37`, `GRCh38`, and `T2T-CHM13`, respectively. We note that without SV harmonization (i.e., without employing `truvari refine`), the average F1 scores were 2 percentage points lower. We note that some tools (e.g., `sawfish`benefit more from SV harmonization, showing up to a 4.9 percentage-point increase in F1 score, whereas some other tools (i.e., `debreak`and `SVision-pro`) do not benefit from it since they do not provide sequence-resolved SV alleles (Supplementary Table 2). We ranked each SV prediction tool according to its performance for each assembly-based SV callset (using the `GRCh38`reference; Figure 3b). We observed shifts in tool rankings depending on the specific ground truth used, even when employing the same reference genome. Moreover, even the choice of using SV harmonization has an impact on the rankings. For instance, we observed that `debreak`, that was one of the best SV caller when not considering SV harmonization, became one of the worst when SV harmonization was taken into account. The reason behind this deterioration must be sought in the inability of the caller in providing sequence resolved SV alleles. Notably, these changes in tool ranking remained dramatic across all other reference genomes used (Supplementary Figure S3 and S4). These results indicate that the performance evaluation of SV detection tools is highly dependent on the ground truth considered, and rankings can drastically change even when the ground truth is originating from the same *de novo* assembly.

**Fig. 3:**
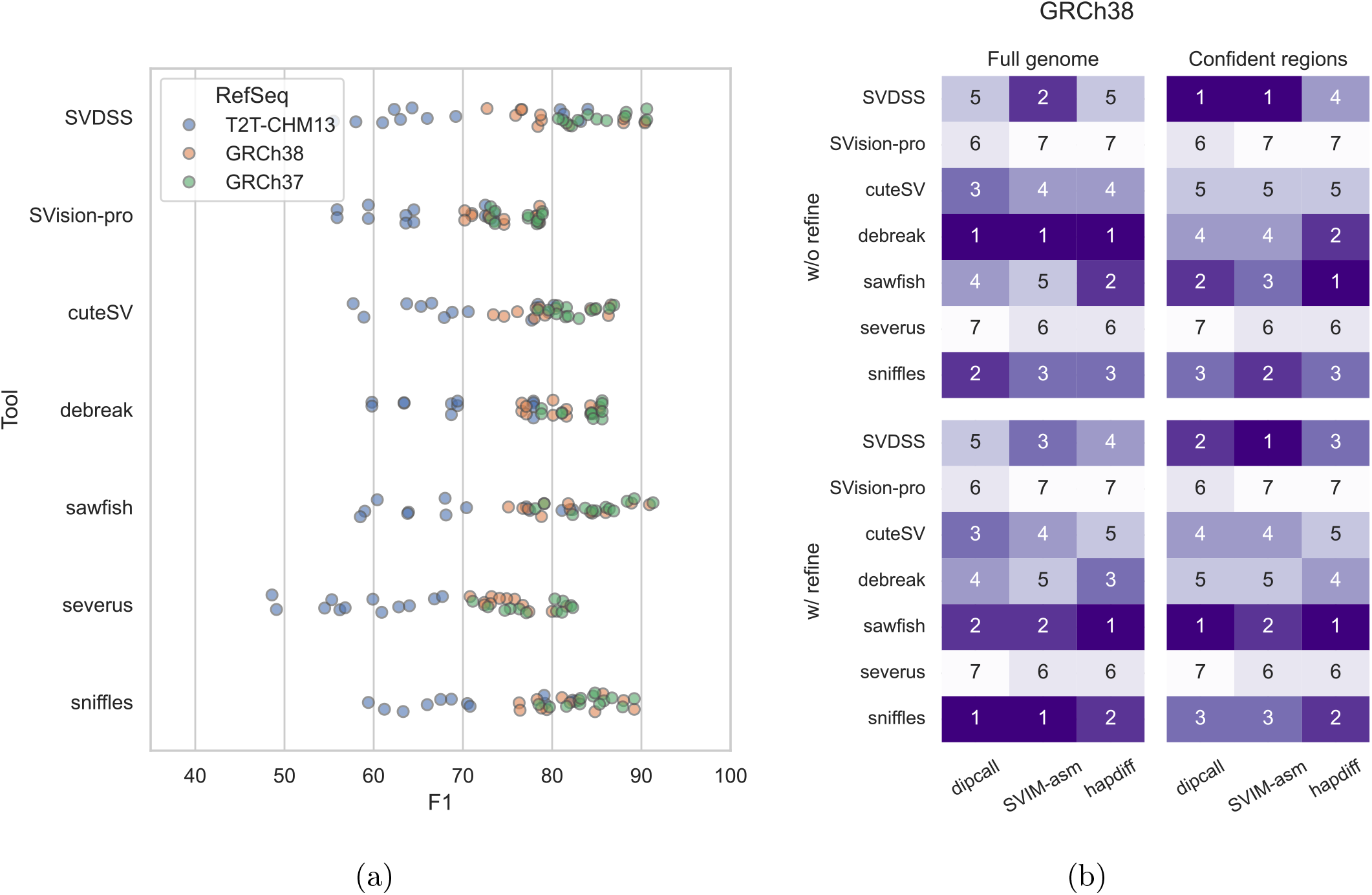
(a) Distribution of F1 score of read-based SV callers when compared to the different ground truth created from HG002 assembly (results stratified by reference sequence only). The full results stratified by reference sequence, assembly-based SV callsets used as ground truth, and `truvari`parameters are presented in Supplementary Table 1. (b) Read-based SV callers ranking with respect to the F1 score reported by `truvari`(without and with SV harmonization) on `GRCh38`(Full genome or Confident regions, as reported by assembly-based callers). Rankings on other references are presented in Supplementary Figures S3 and S4).

We next stratified the results using the v3.6 genome stratification provided by the GIAB project [34], focusing on “all difficult regions” and “not in all difficult regions” (Figure 4). In the “easy” regions of the genome (i.e., those classified as “not in all difficult regions”), all tools achieved very high accuracy across all three reference genomes, particularly when considering as ground truth the assembly-based SV callsets generated with `SVIM-asm`or `hapdiff`(average F1 score of 96%). However, when using the assembly-based SV callset generated by `dipcall`as ground truth, F1 score of all tools decreased to 94%. This slightly reduction of accuracy further highlights the discrepancies in SV evaluation that arise from the choice of ground truth, even in those regions of the genome considered not challenging. Moreover, all tools achieved drastically lower accuracy when the analysis is restricted to hard regions of the genome (Figure 4). On `GRCh37`and `GRCh38`, the average F1 score on hard regions drops to 77% and 73%, respectively. The drop in F1 score between easy and hard regions is even more pronounced on the `T2T-CHM13`reference genome where the average F1 score is 56% and the highest F1 score is 70.2%, providing more evidence of the complexity of accurate SV calling in repetitive regions of the genome that are unique to the recent `T2T-CHM13`reference genome (Figure 4). Results obtained without using SV harmonization are presented in Supplementary Figure S5. As in our previous analysis, SV harmonization increased the F1 scores of most callers, with an average increase of 0.89 in easy regions and 1.44 in hard regions of the genome. Full results on GIAB stratification are provided in Supplementary Tables 3 and 4.

**Fig. 4:**
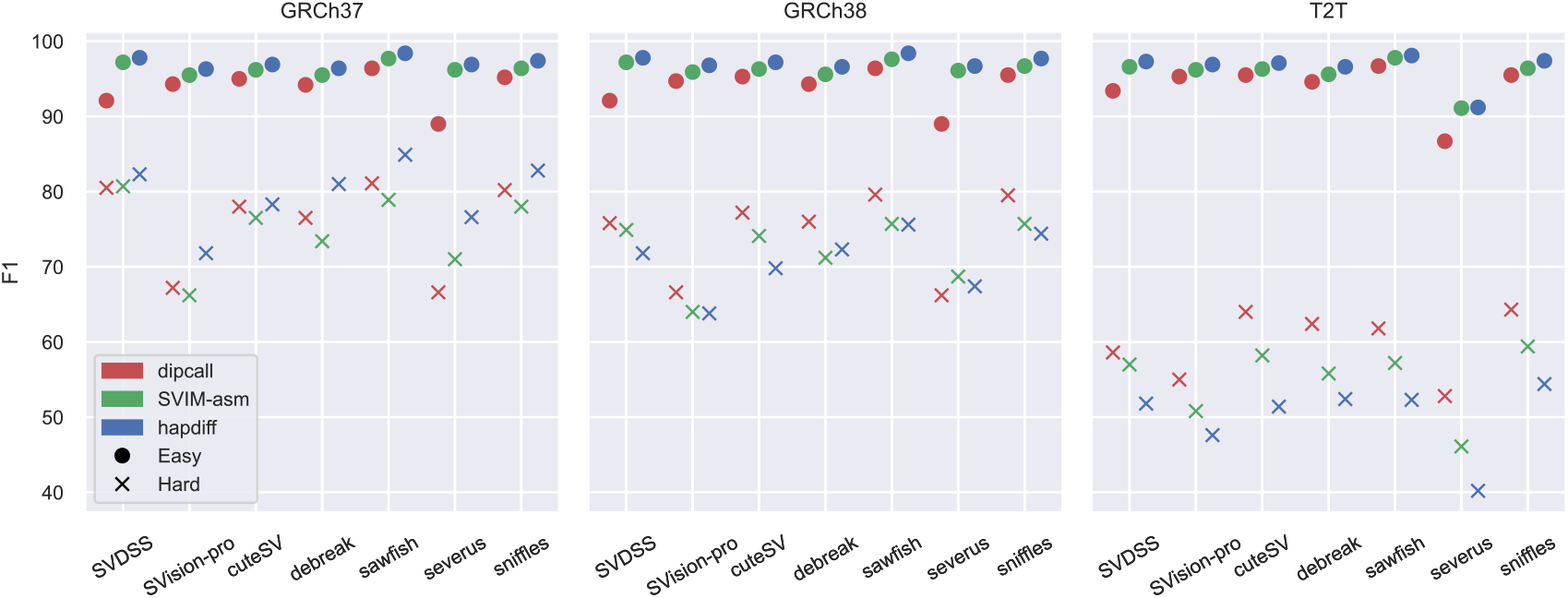
F1-measure of read-based callers. Results are stratified by reference genome, ground truth (computed with assembly-based SV callers), and GIAB v3.6 genome stratification (“Easy” refers to the regions tagged as “not in all difficult regions” whereas “Hard” to those tagged as “all difficult regions” in the stratification files). F1 scores obtained with SV harmonization, i.e., via `truvari refine`. Results without harmonization are presented in Supplementary Figure S5).

### Benchmarking of read-based SV callsets against curated ground truths

Finally, we examined the impact of using two different curated SV callsets published by the Genome in a Bottle (GIAB) consortium for the HG002 sample: the GIAB v0.6 callset [18] and the recent GIAB v1.1 callset^9^ (preliminary draft, currently under investigation). We focused our main investigation on the confident regions provided by the consortium and the accuracy results obtained after SV harmonization, i.e., using `truvari refine`(Figure 5). When comparing the two GIAB curated SV callsets, we observed substantial changes in the ranking of tool performance depending on which version was used. On the older v0.6 curated SV callset, all read-based approaches achieved very high recall (>95%) and good precision, with F1 scores ranging from 92.7% to 96.4%. Accuracy of all tools drops consistently when considering the more recent v1.1 curated SV callset. All tools achieved similar precision (>90%) but lower recall (<87%). The lowest accuracy is achieved by all tools when evaluated on the v1.1 curated SV callsets from the `T2T-CHM13`reference genome. We, indeed, observed a considerable drop in the overall performance of all tools (based on F1 score) when evaluated against the more complete curated SV callset (GIAB v1.1, `T2T-CHM13`reference), with average F1 score decreasing from 95.7% (GIAB v0.6, `GRCh37`reference) to 78.8% (Figure 5). This result further indicates that the performance evaluation between methods is highly dependent on the ground truth employed and modifications to these sets can seriously alter the observed performance hierarchy. The performance drop is even more pronounced when considering the full genome, thus not restricting the evaluation to those regions of the genome identified as high-confidence by the consortium. Supplementary Figure S6 reports the results of this analysis on full genome. We notice that when considering the more recent v1.1 curated callset and the more complete `T2T-CHM13`reference sequence, the F1 score of all tools drop to around 60% which is comparable to the results obtained when evaluating against assembly-based SV callsets (Figure 3a). This further indicates that even state-of-the-art methods for SV detection using long-read WGS data still struggle in some of the more complex regions of the genome.

**Fig. 5:**
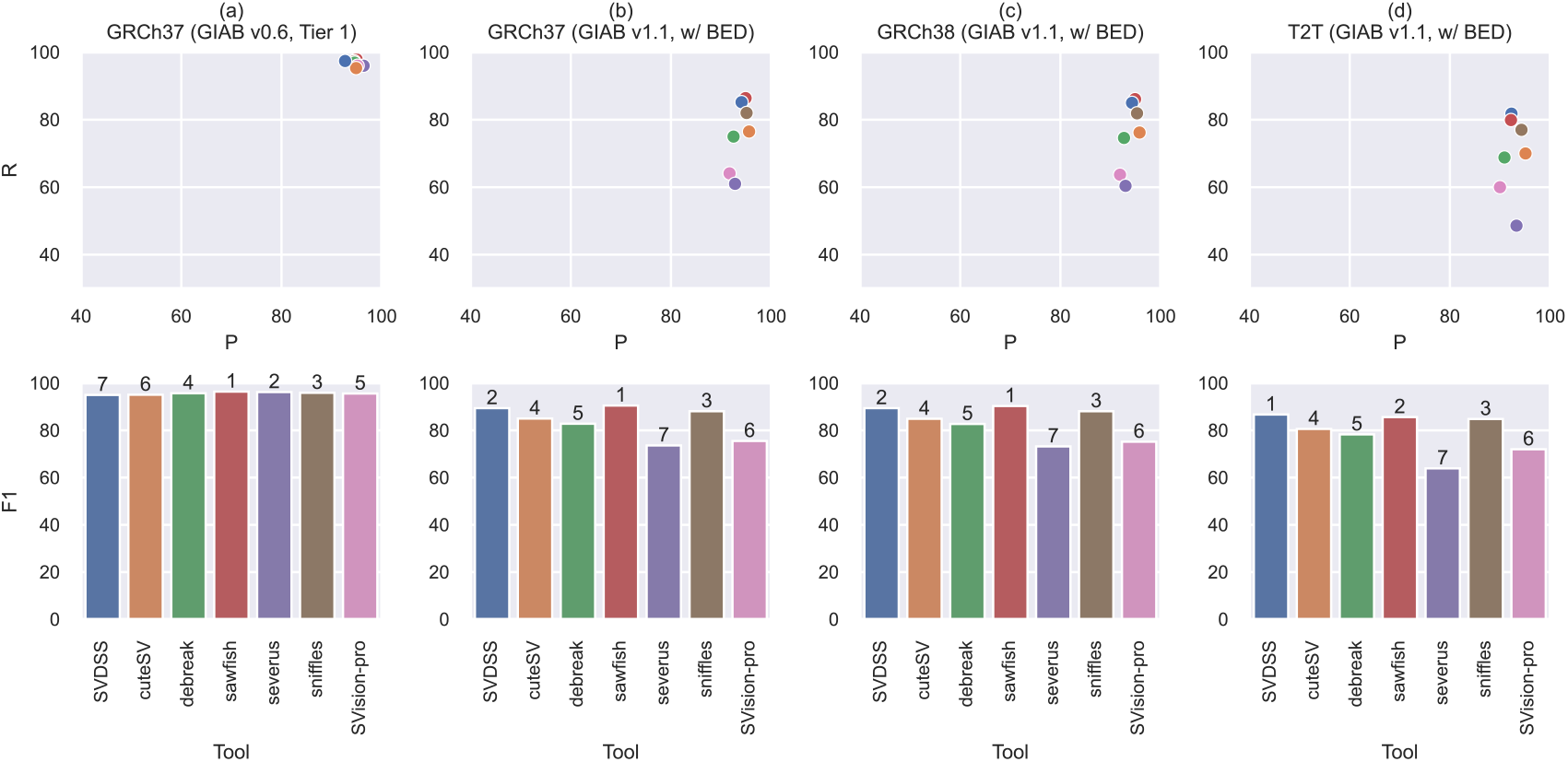
Performance evaluation against GIAB gold standard SV callsets. Precision (P), Recall (R), and F1 score (F1) of different methods evaluated using different curated SV callsets as ground truths and reference genomes (which are noted in the top of every column). The number on top of each bar (second row) is the rank of each method. Note: `SVision-pro`results are not included due to `truvari`requiring sequence-level comparison.

Even in this comparison with curated ground truths, SV harmonization allowed to marginally increase the accuracy of each tool, with an increase of 2 percentage points on average. However, also in this evaluation, we observed that some tools (e.g., `sniffles2`and `sawfish`) benefited more from this normalization, with an increase of up to 6.40 percentage points. Results without SV harmonization are presented in Supplementary Figure S6. Full results of this analysis are provided in Supplementary Tables 5 and 6.

## 3 Methods

To streamline the SV evaluation task, we have devised a modular and extensible framework for the analysis and comparison of SV callsets produced by both long-reads and assembly-based callers. The framework is provided as a Snakemake pipeline and is freely available at https://github.com/ldenti/svbench-fw, along with all the scripts used to analyze the callsets. Our framework can be easily used to reproduce the results described in this paper and also to replicate the analyses on new individuals. The input required to run the framework analyzes consists of: the selected reference genome, a long-read WGS sample, a diploid assembly of the individual under investigation, and genomic annotations (e.g., tandem repeats annotation and genome stratifications). The framework includes the 8 SV callers presented in Table 1 and 3 assembly-based approaches, namely `dipcall`(v0.3), `SVIM-asm`(v1.0.3), and `hapdiff`(commit `e0abbb9`), but can be easily extended with additional methods. Whenever a new method for SV calling (from long reads or assembly) is proposed, it can be included in our framework by simply adding a new rule to the `Snakemake`pipeline.

### Assembly-based callers

The 3 assembly-based callers are run using their default parameters and setting the minimum SV size to 30bp (for `SVIM-asm`). Since `dipcall`outputs SNPs and indels, its callset is post-processed with `bcftools view`to include only structural variants longer than 30bp. In addition, since `dipcall`outputs multiallelic variants that are not supported by `truvari`, these calls are split using `bcftools norm`. We notice that we decided to consider structural variants longer than 30bp since this is the default value used by `truvari`when parsing the truth set. However, since structural variants are typically defined as alterations longer than 50bp, when comparing the three assembly-based SV callsets, we considered only variants longer than 50bp.

### Reads alignment and short variants calling

The input HiFi reads are aligned with `minimap2`(v2.28) using the following options: `-ax map-hifi –MD –eqx -Y`. Since `Severus`requires as input a phased VCF, to obtain the required files we relied on `deepvariant`(v1.8.0) and `whatshap`(v2.5), both run with default parameters.

### Long reads-based callers

All long-reads based callers are run using their default parameters or setting the same value for common parameters. The minimum mapping quality for read alignments is set to 20, the minimum size for calls is set to 50bp, and the minimum support for a call to 4 (when the tool does not provide an automatic way to detect this). In more detail, `cuteSV`(v2.1.2) is run using `--min_support 4`, `--min_size 50`, and `--max_cluster_bias_INS 1000 --diff_ratio_merging_INS 0.9 --max_cluster_bias_DEL 1000 --diff_ratio_merging_DEL 0.5`, as suggested by the authors. We note that we did not change the `--min_mapq`option since it is by default set to 20. `SVDSS`(v1.0.5) is run setting `--min-cluster-weight 4`. `sniffles2`(v2.3) is run using all default parameters and providing a tandem repeat annotation BED file (downloaded from the `pbsv`repository) via the `--tandem-repeats`option. Since `sniffles2`estimates the minimum support automatically based on coverage, we did not change the value of `--minsupport`option. `Severus`(v1.4.0) is run using `--min-mapq 20 --min-support 4`and providing the tandem repeats annotation via the `--vntr-bed`option. `debreak`(v1.3) is run using `--min_size 50 --min_quality 20`and setting the aligner with `--aligner minimap2`. Since `debreak`estimates the minimum support automatically from alignment depth, we let it decide the best value for this parameter.

`SVision-pro`(v2.4) is run in germline mode (`--detect_mode germline`) using the neural network model *model_liteunet_256_8_16_32_32_32.pth* (set via the `--model_path`option). Additionally, we set `--min_supp 4`and `--preset hifi`. `sawfish`(v2.0.0) is run using `--min-indel-size 50`and `--min-sv-mapq 20`.

*SV callsets evaluation* To compare SV callsets, we used `truvari`(v5.3.0) and tested two distinct parameter settings:

1. `truvari-def`: default `truvari`parameters, with the exceptions of `--passonly`, `--pick ac`, and `--dup-to-ins`to accommodate specific tool outputs.
2. `truvari-wbed`: same as `truvari-def`on only a subset of genomic regions, considered as *“confident”* regions (we used the `--includebed`option to pass the regions BED file)

Since running SV harmonization through `truvari refine`requires the input SV callsets to be phased, if a caller does not provide phased genotypes, we genotyped the variants using `hiphase`(v1.5.0) ran with default parameters. We ran SV harmonization with the command:

~~~
truvari refine --reference {reference.fa} \
--regions {truvari_directory}/candidate.refine.bed \
--coords R \
--use-original-vcfs \
--threads {threads} \
--align mafft \
{truvari_directory}
~~~

### 3.1 Data

In our experimental evaluation, we considered the HG002 individual, since it is one of the most well studied individuals in terms of structural variants and many benchmark callsets are available in the literature (e.g., GIAB v0.6 and GIAB v1.1). In our analysis, we considered the HG002 data provided by the Human Pangenome Reference Consortium [31]. The HG002 contigs were extracted using `agc`[35] from the full set of assemblies provided by the consortium (year 1 freeze^10^) whereas the PacBio HiFi sample (Sequel II System with Chemistry 2.0, 15kb Library, 36x coverage) was downloaded from the HPRC s3 server^11^ (runs `m64012_190920_173625`, `m64012_190921_234837`, `m64015_190920_185703`, and ftp://ftp-trace.ncbi.nih.gov/1000genomes/ftp/technical/reference/m64015_190922_010918). In all our analysis, we considered three reference sequences, namely `GRCh37`(from ftp://ftp.ncbi.nlm.nih.gov/genomes/all/GCA/human_g1k_v37.fasta.gz), `GRCh38`(from 000/001/405/GCA_000001405.15_GRCh38/seqs_for_alignment_pipelines.ucsc_ids/GCA_ 000001405.15_GRCh38_no_alt_analysis_set.fna.gz), and `T2T-CHM13`(from https://s3-us-west-2.amazonaws.com/human-pangenomics/T2T/CHM13/assemblies/analysis_set/chm13v2.0.fa.gz).

## 4 Discussion

We identified critical and often unexpected divergence in the performance evaluations of different SV calling methods, largely driven by commonly applied user-defined choices in analysis workflows. We believe this inconsistency in SV calling evaluation is partly attributable to variations in how ground truths are constructed (or selected), as well as differences in the reference genomes used during benchmarking. These discrepancies highlight the critical need for standardization in SV evaluation, particularly when benchmarking SV calling methods or assessing the overall accuracy of SV prediction approaches for downstream analyses.

To better understand the variability introduced by assembly-based SV callset generation methods (which are typically used to create the ground truth), we manually examined several loci on the `T2T-CHM13`reference genome where SV calls from three such methods, namely `SVIM-asm`, `dipcall`, and `hapdiff`, disagreed, despite all being derived from the same *de novo* assembly. Our investigation revealed two major factors contributing to discrepancies between these truth sets. First, the methods differ in how they represent haplotype-resolved variants. The `SVIM-asm`and `hapdiff`approaches tend to report homozygous alternative alleles (1|1), while `dipcall`more often outputs two distinct heterozygous variants, sometimes fragmenting SVs into separate events on each haplotype (Supplementary Figure S7 reports an example). For instance, in some loci, `dipcall`reported two nearby insertions differing by a single base, whereas `SVIM-asm`merged them into a single call. Second, differences in alignment strategy play a key role. `hapdiff`uses custom `minimap2`parameters optimized for long SVs, while `dipcall`uses default settings. Since `SVIM-asm`was run on `dipcall`alignments in our study, discrepancies between `hapdiff`and the others are likely due to alignment differences. Supplementary Figures S8 and S9 reports two loci where the alignments used by `hapdiff`, differently from the ones produced by `dipcall`, exhibit no SV signature on the haplotypes. Together, these findings highlight the critical impact of methodological choices in SV ground truth generation, ranging from alignment strategies to variant representation and merging heuristics. Such differences complicate the SV ground truth construction and emphasize the need for standardized, transparent protocols for the benchmarking of SV callers in genomic analyses. Similarly, we observed substantial changes in the accuracy rankings of SV calling methods when evaluated using different assembly-based truth sets or different curated SV callsets, such as GIAB v0.6 versus GIAB v1.1. This highlights that the performance of SV callers is not readily transferable between ground truths without careful consideration of their limitations and subtle differences. Equally importantly, we observed a decline in the performance of all SV callers when using the `T2T-CHM13`reference genome, highlighting the limitations of current methods in accurately detecting SVs within more complex genomic regions accessible only through `T2T-CHM13`reference genome, even when leveraging long-read WGS data.

To address the observed discrepancies, we developed a pipeline, freely available at https://github.com/ldenti/svbench-fw, for the comprehensive construction of multiple ground truths using state-of-the-art assembly-based SV detection methods, all derived from same high-quality *de novo* assemblies. This pipeline incorporates several state-ofthe-art assemblyand long reads-based SV calling methods. Benchmarking old and new SV calling methods against a diverse collection of truth sets enables a more robust and comprehensive evaluation of SV calling performance.

Finally, we note that this study did not assess the impact of varying parameters or configurations for each method included in the evaluation. Instead, we used the default settings recommended by the developers of each tool. Parameter tuning introduces an additional layer of complexity that could substantially affect performance and lead to an even broader range of possible relative rankings, further complicating any direct comparisons.

## Acknowledgements

The authors thank Heng Li for his valuable feedback and suggestions.

This research work has received funding from the European Union’s Horizon programme under the Horizon Europe grant agreement (ASVA-CGR No. 101180581 to L.D.). This work has also been supported in part by National Science Foundation (NSF) award DBI-2042518 to F.H. This research was supported in part by the Slovak Grant Agency VEGA grants 1/0538/22 (T.V.) and 1/0140/25 (B.B.). P.B. has received funding from the European Union’s Horizon 2020 research and innovation programme under the Marie Skłodowska-Curie grant agreement PANGAIA No. 872539 and ITN ALPACA N.956229 and from Next Generation EU - Mission 4, MIUR 2022YRB97K, PINC, Pangenome Informatics: from Theory to Applications.

## Footnotes

9 https://ftp-trace.ncbi.nlm.nih.gov/ReferenceSamples/giab/data/AshkenazimTrio/analysis/NIST_HG002_DraftBenchmark_defrabbV0.019-20241113/

10 https://zenodo.org/record/5826274/files/HPRC-yr1.agc

11 https://s3-us-west-2.amazonaws.com/human-pangenomics/index.html?prefix=working/HPRC_PLUS/HG002/raw_data/P

