## Supplementary Figures for "Anyone can be the best: Impact of diverse methodologies on the evaluation of structural variant callers"

*Luca Denti, Thomas Krannich, Tomas Vinar, Rayan Chikhi, Paola Bonizzoni, Brona Brejova, and Fereydoun Hormozdiari*

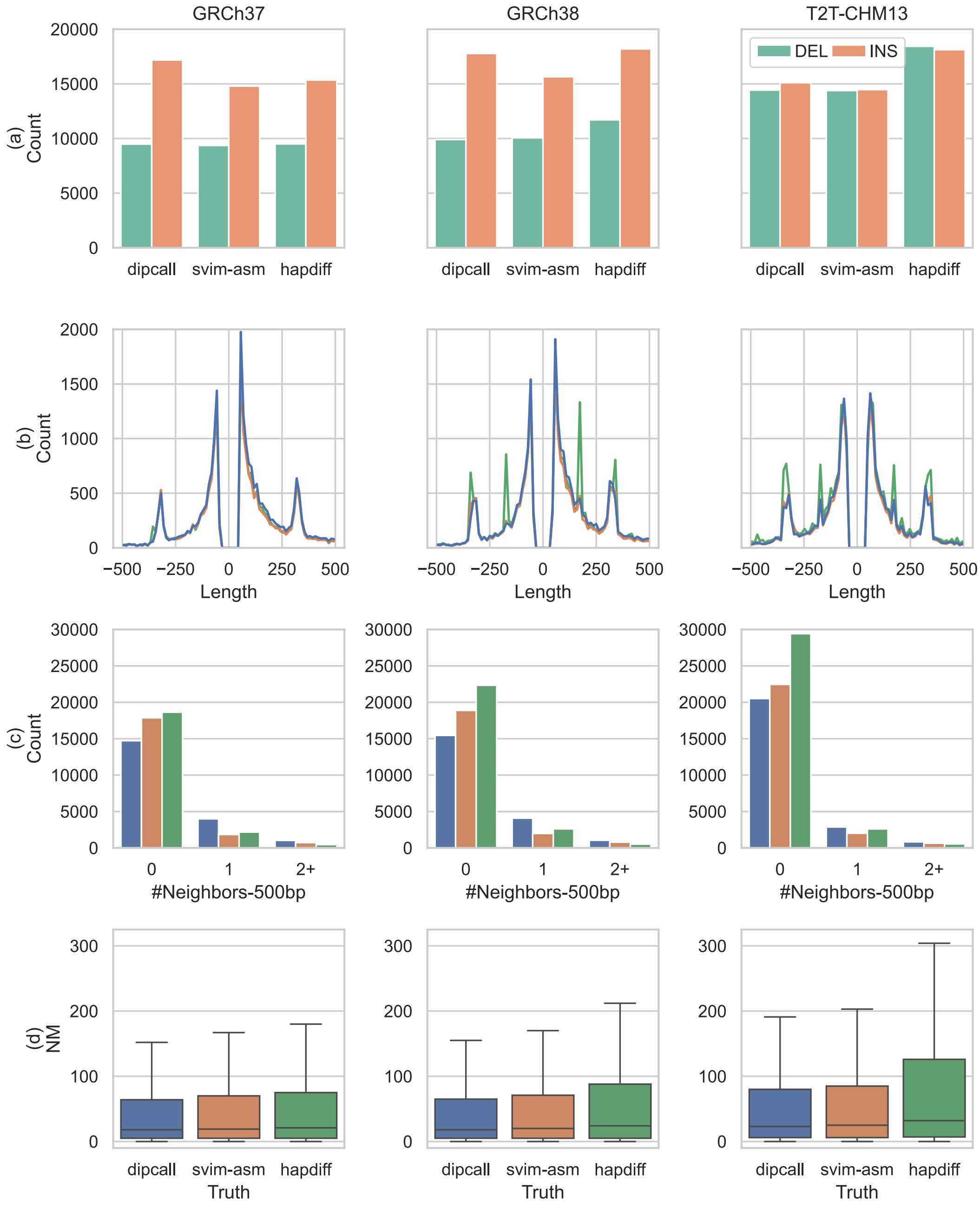


Fig. S1: Comparison of state-of-the-art assembly-based SV callers using the same *de novo* assembly against three reference genomes (full genome results). (a) Number of SVs reported for each approach based on their type (INSertions and DELetions). (b) Length distribution of unbalanced structural variants (negative length for deletions). (c) Number of neighboring SVs within 500bp. (d) Distribution of the total number of mismatches and gaps (`NM` field) when aligning "alternative" contigs created from the VCF produced by each method against the true contigs.

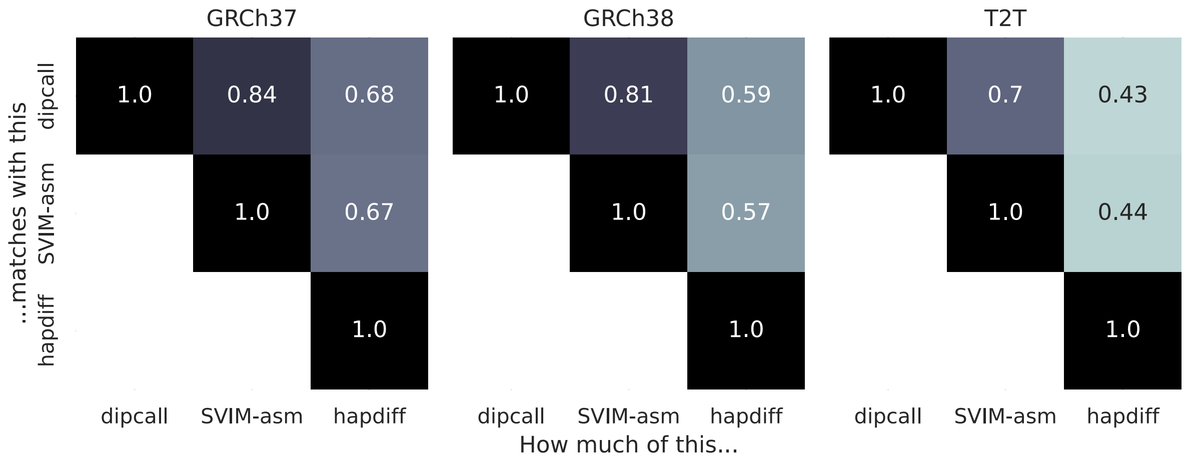


(a) Full genome, without SV harmonization

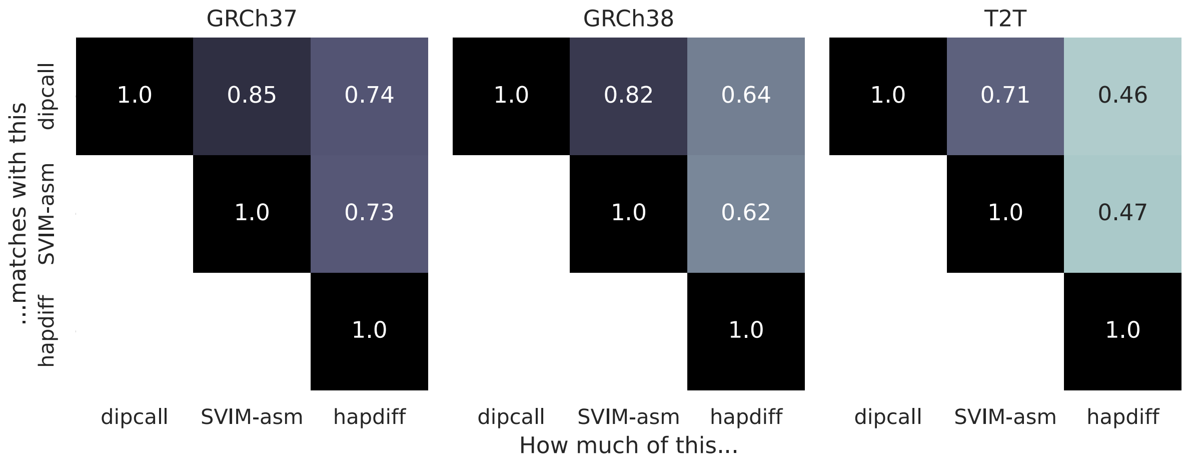


(b) Full genome, with SV harmonization

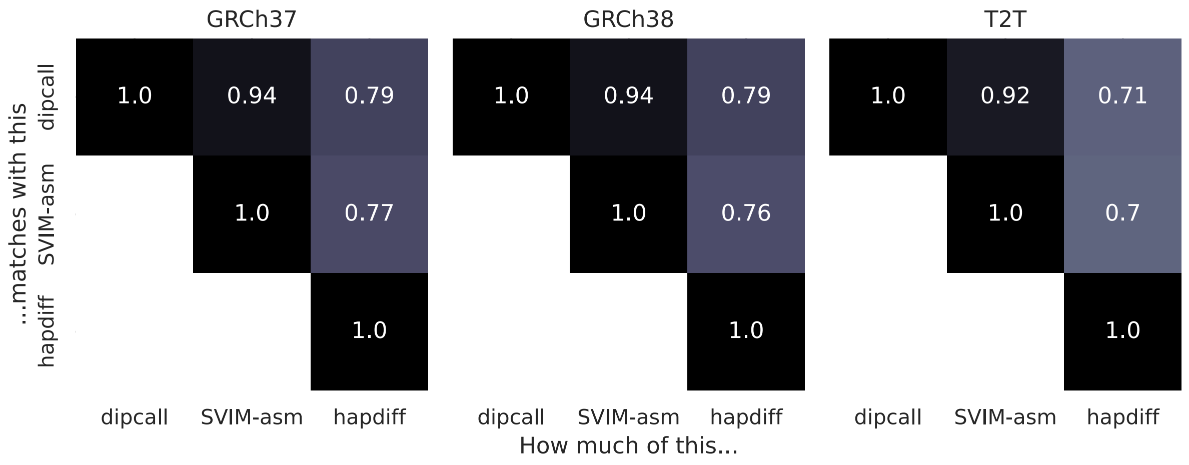


(c) Confident regions of the genome, without SV harmonization

Fig. S2: Pairways similarity between assembly-based SV callers. Each cell presents the accuracy (TP/(TP+FP+FN)) of the column tool (considered as the callset) w.r.t. the row tool (considered as the truth set).

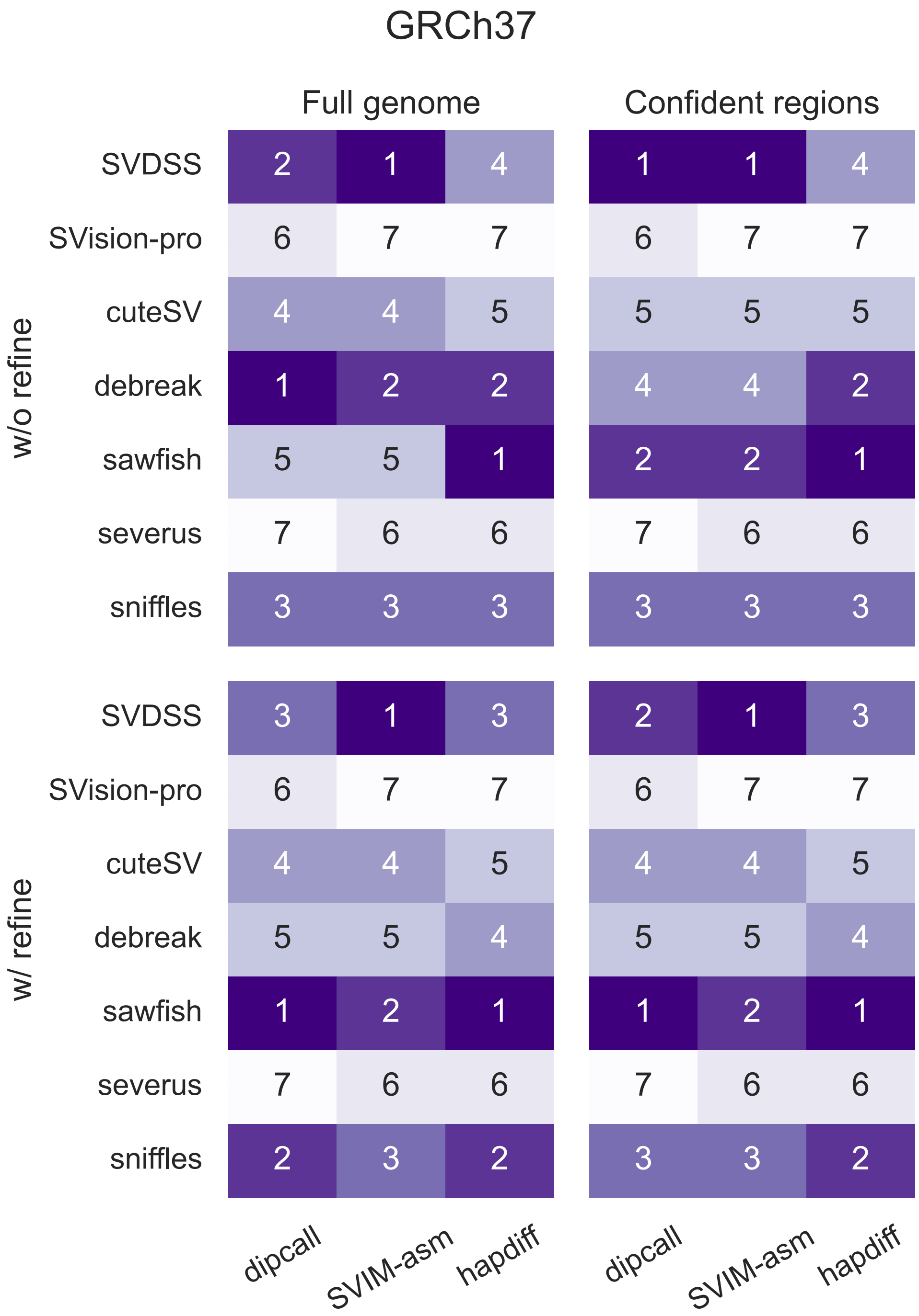


Fig. S3: Read-based SV callers ranking w.r.t. F1 score reported by truvari (hg19).

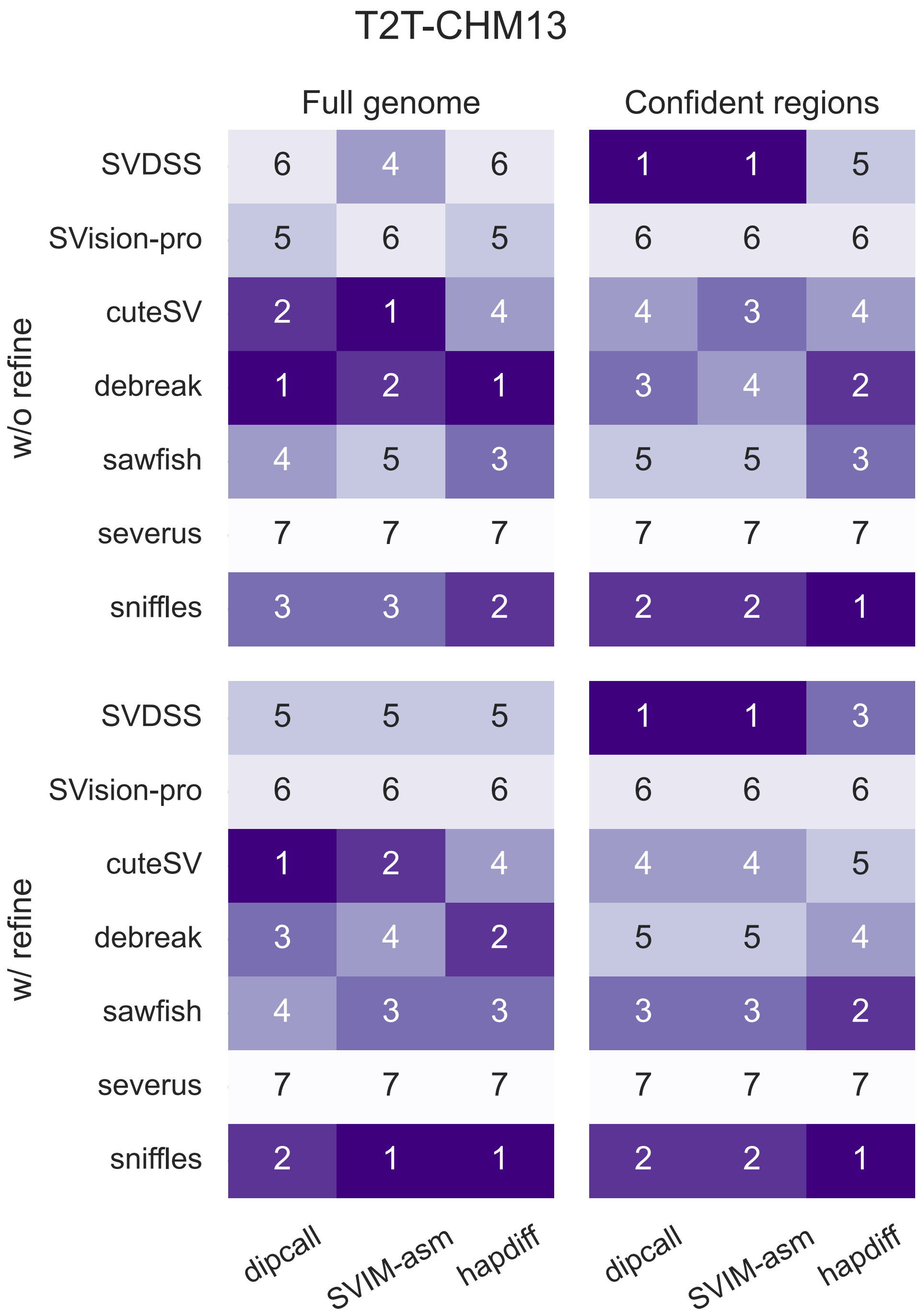


Fig. S4: Read-based SV callers ranking w.r.t. F1 score reported by truvari (t2t).

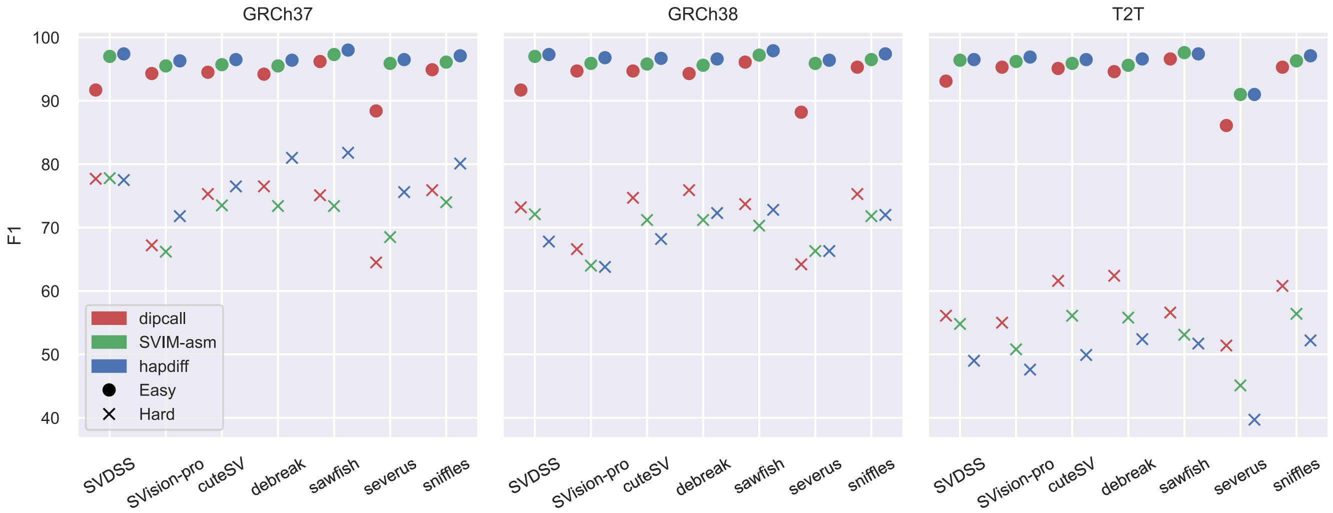


Fig. S5: F1-measure of read-based callers. Results are stratified by reference genome, ground truth (computed with assembly-based SV caller), and GIAB v3.6 genome stratification ("Easy" refers to the regions tagged as "not in all difficult regions" whereas "Hard" to those tagged as "all difficult regions" in the stratification files). F1 scores obtained without SV harmonization (i.e., via `truvari bench`).

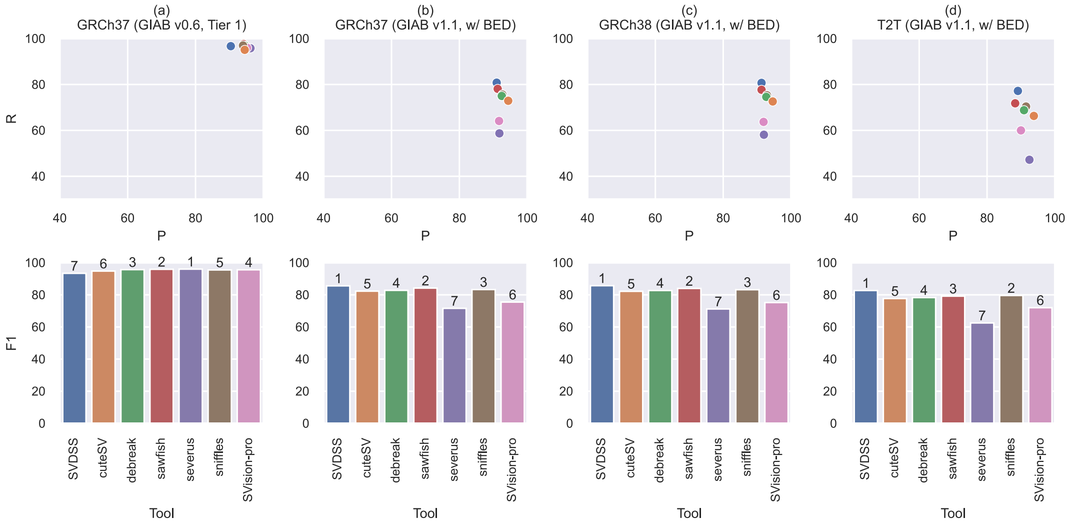


(a) Confident regions of the genome, without SV harmonization

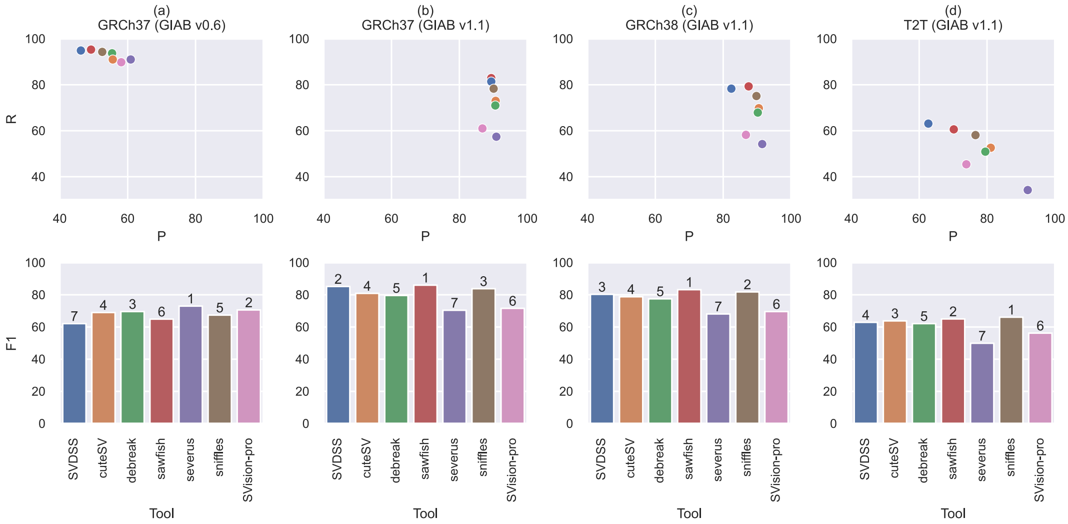


(b) Full genome, with SV harmonization

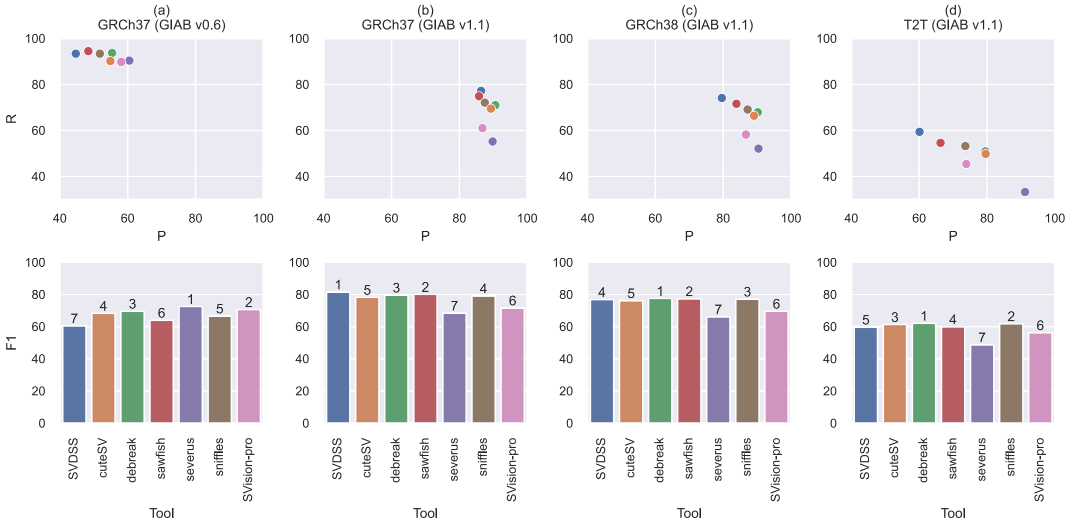


(c) Full genome, wihtout SV harmonization

Fig. S6: Performance evaluation against GIAB curated SV callsets. Precision (P), Recall (R), and F1 score (F1) of different methods evaluated using different truth sets and reference genomes (which are noted in the top of every column). The number on top of each bar (second row) is the rank of each method.

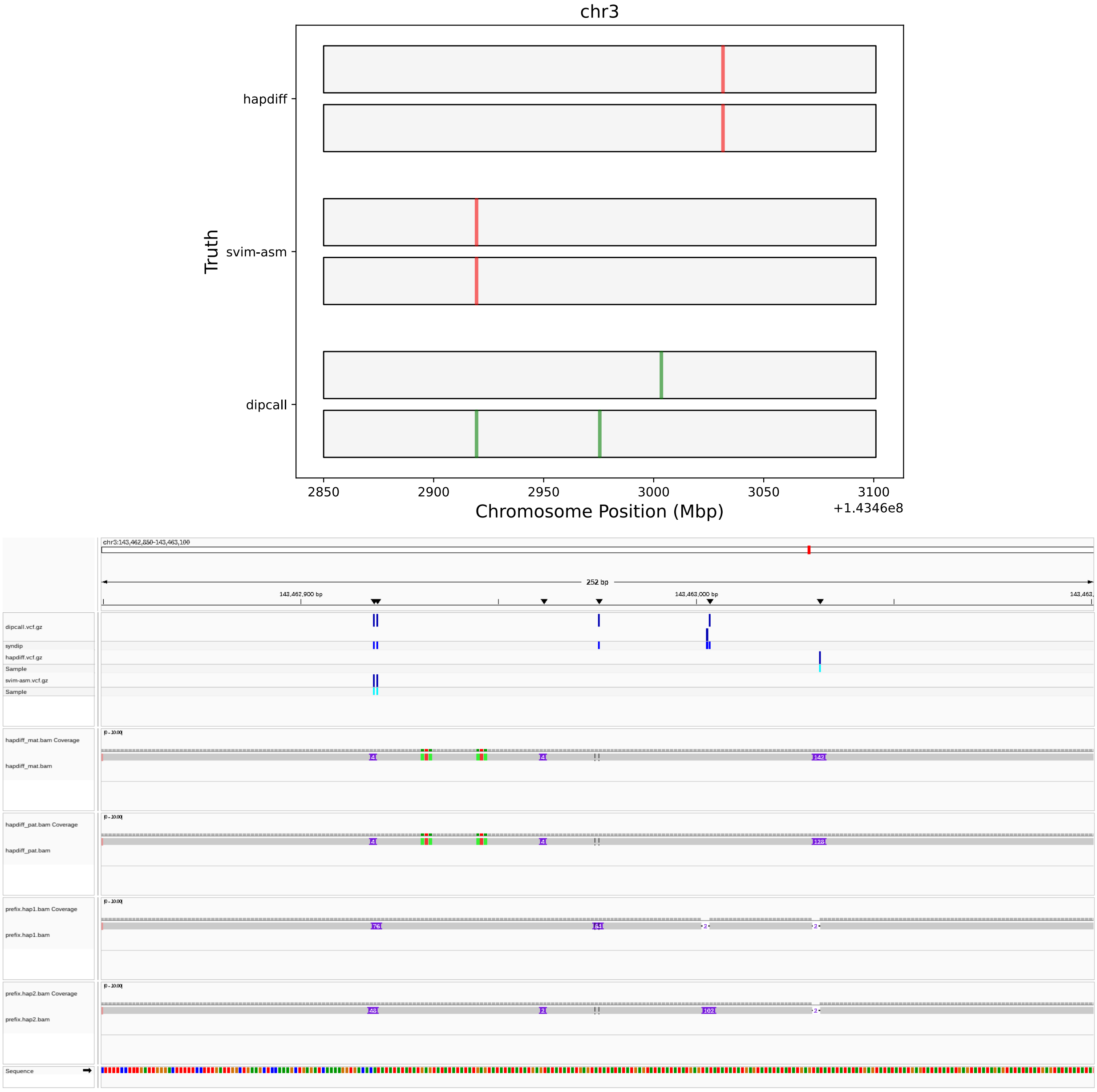


Fig. S7: Locus `chr3:143462850-143463100` (`T2T-CHM13` reference genome). Top panel: schematic representation of the structural variants reported by the three assembly-based callers. Red bars represent homozygous calls (`1|1`) while green bars represent heterozygous calls (`0|1` and `1|0`). Bottom panel: IGV screenshot of this locus. The first three tracks are the VCF produced by the assembly-based callers. Fourth and fifth tracks (*hapdiff_ mat.bam* and *hapdiff_ pat.bam*) are the alignments computed and used by `hapdiff`. Bottom tracks (*prefix.hap1.bam* and *prefix.hap2.bam*) are the alignments computed by `dipcall` (and used by `dipcall` and `SVIM-asm`).

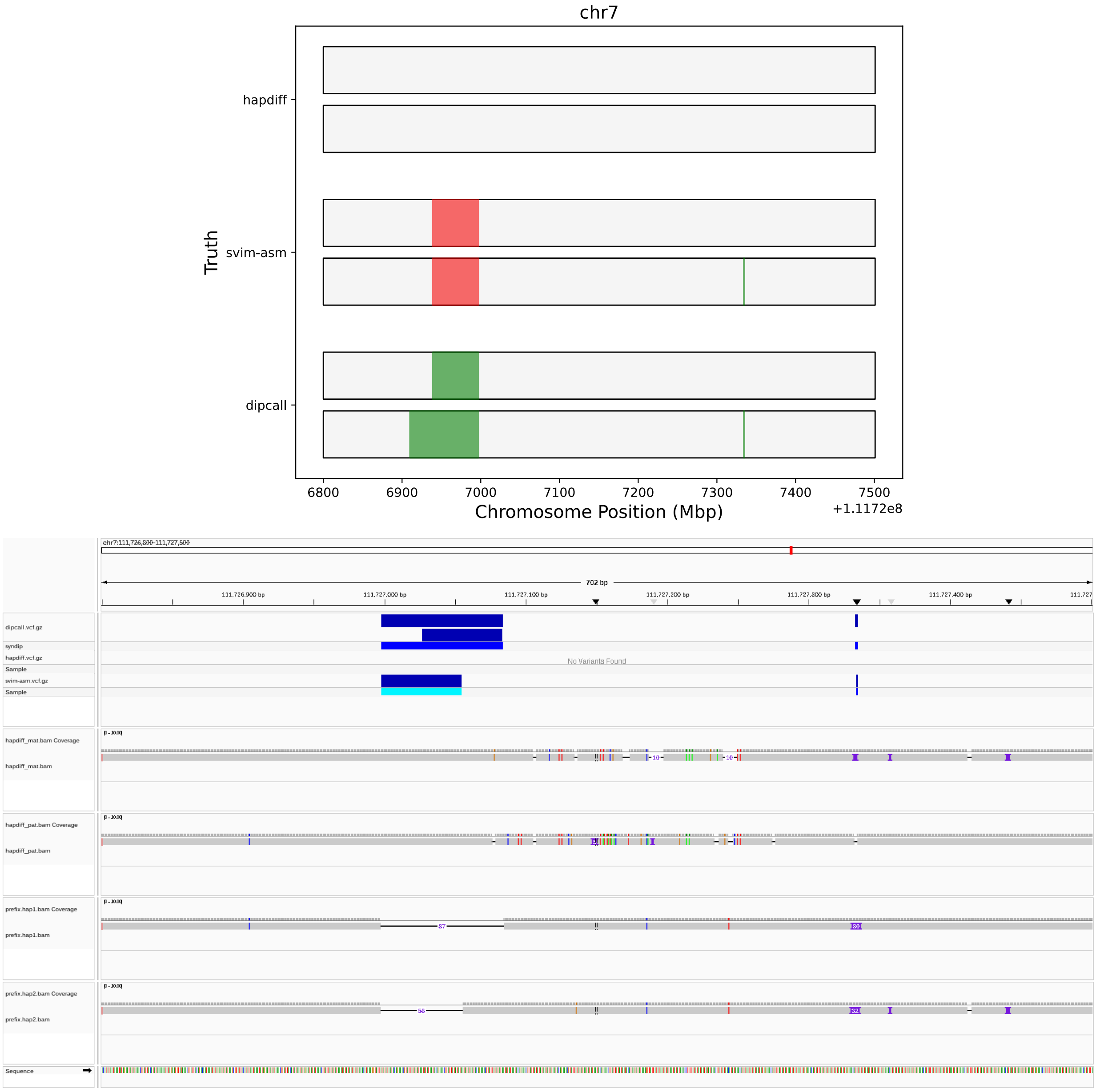


Fig. S8: Locus `chr7:111726800-111727500` (`T2T-CHM13` reference genome). Top panel: schematic representation of the structural variants reported by the three assembly-based callers. Red bars represent homozygous calls (`1|1`) while green bars represent heterozygous calls (`0|1` and `1|0`). Bottom panel: IGV screenshot of this locus. The first three tracks are the VCF produced by the assembly-based callers. Fourth and fifth tracks (*hapdiff_mat.bam* and *hapdiff_pat.bam*) are the alignments computed and used by `hapdiff`. Bottom tracks (*prefix.hap1.bam* and *prefix.hap2.bam*) are the alignments computed by `dipcall` (and used by `dipcall` and `SVIM-asm`).

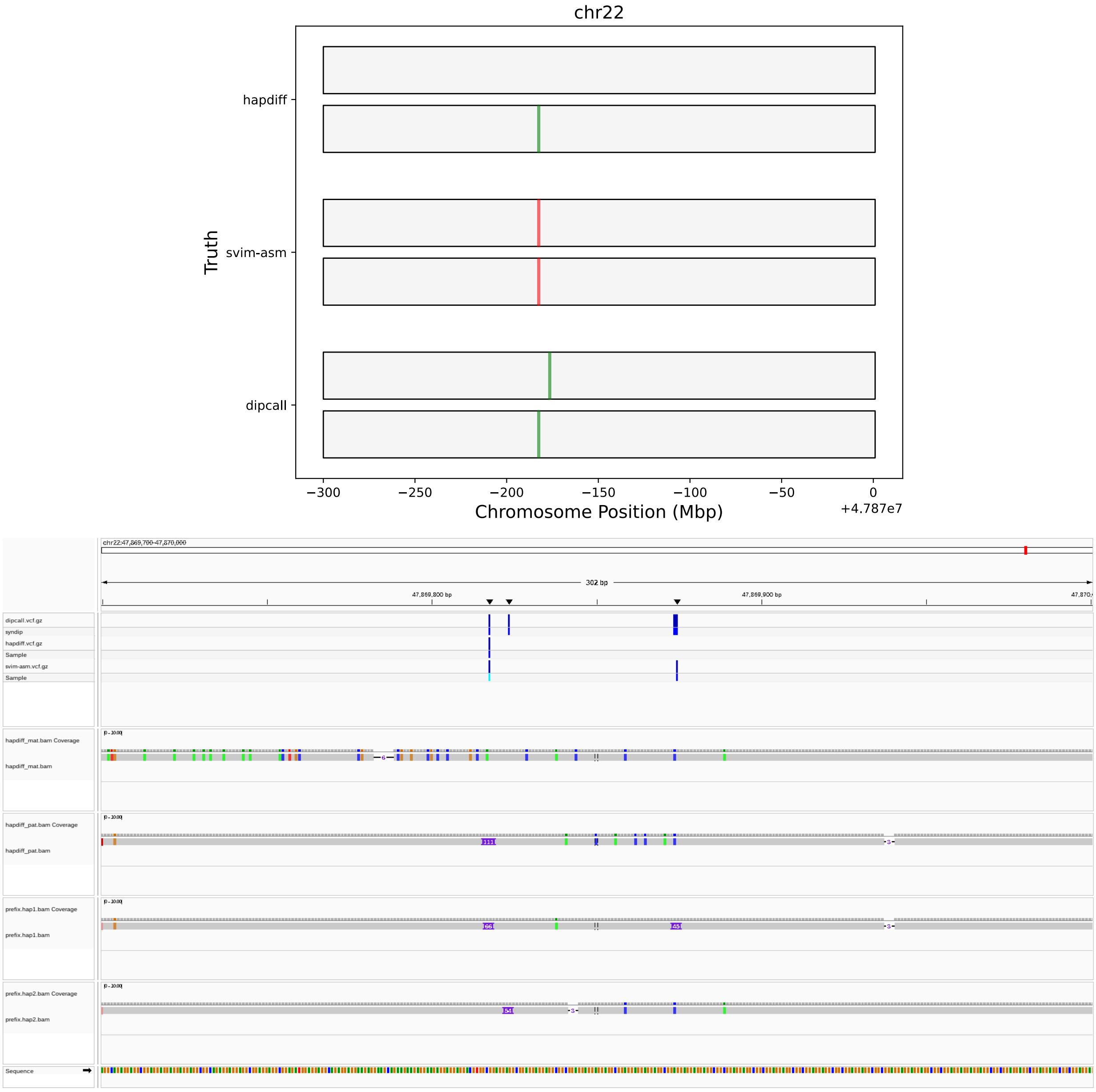


Fig. S9: Locus `chr22:47869700-47870000` (`T2T-CHM13` reference genome). Top panel: schematic representation of the structural variants reported by the three assembly-based callers. Red bars represent homozygous calls (`1|1`) while green bars represent heterozygous calls (`0|1` and `1|0`). Bottom panel: IGV screenshot of this locus. The first three tracks are the VCF produced by the assembly-based callers. Fourth and fifth tracks (*hapdiff_mat.bam* and *hapdiff_pat.bam*) are the alignments computed and used by **`hapdiff`**. Bottom tracks (*prefix.hap1.bam* and *prefix.hap2.bam*) are the alignments computed by `dipcall` (and used by `dipcall` and `SVIM-asm`).
